# Optimal biochemical information processing at criticality

**DOI:** 10.1101/543348

**Authors:** Angel Stanoev, Akhilesh P. Nandan, Aneta Koseska

## Abstract

How cells utilize surface receptors for chemoreception is a recurrent question spanning between physics and biology over the past few decades. However, the dynamical mechanism for processing time-varying signals is still unclear. Using dynamical systems formalism to describe criticality in non-equilibrium systems, we propose generic principle for temporal information processing through phase-space trajectories using dynamic transient memory. In contrast to short-term memory, dynamic memory generated via ghost attractor enables signal integration depending on stimulus history, and thus balance between stability and plasticity in receptor responses. We propose that self-organization at criticality can arise through fluctuation-sensing mechanism, illustrated for the experimentally established epidermal growth factor sensing system. This framework applies irrespective of the intrinsic node dynamics or network size, as we show using also a basic neuronal model. Processing of non-stationary signals, a feature previously attributed only to neuronal networks, thus uniquely emerges for biochemical networks organized at criticality.

## I. INTRODUCTION

In a wide variety of biological processes including embryogenesis, immune cells motility, wound healing or cancer metastasis [1–3], cells sense and respond to time-varying chemical signals that reflect the non-stationary environment to which they readily adapt. Sensing of chemical signals occurs through receptor-coupled enzymatic activity that transmits the information about the environment through chemical modifications. The receptor activity dynamics on the other hand emerges from the biochemical network in which the receptor is embedded. To perceive and process continuous streams of multimodal inputs, the cellular sensing system must satisfy two general, but seemingly opposed demands. Sensing requires plasticity in the receptor activity responses to enable rapid responsiveness to continuous changes in the environment, whereas robust signaling responses require prolonged receptor activity through a transient memory to be maintained on the cell surface after signal removal [4]. Such a memory is necessary to process the temporal dependencies that are inherent in time-varying signals [5]. However, a theoretical framework that explains how these opposed features emerge on the level of single cells has been missing, since common models for computation such as Turing machines or attractor networks cannot capture this dynamics. While Turing machines describe off-line computation on (static) discrete inputs [6], attractor networks that use multiistability for memory realization settle into a fixed-point and can thereby temporally limit the perception of upcoming stimuli [4].

Processing of time-varying chemical signals by cell surface receptors on the other hand dynamically resembles the real-time computations of sensory stimuli carried out by neural microcircuits in the cerebral cortex [7, 8]. To overcome the limitations of Turing computation in the latter case, universal frameworks using transient dynamics and state-dependent trajectories have been proposed as principal forms of computations that rely on the high-dimensionality of the networks with complex intrinsic neuronal dynamics on several temporal scales [7, 9, 10]. These formalisms however cannot be directly translated to the equivalent computational problem in biochemical networks. First, receptors do not exhibit intrinsic complex activity dynamics such as neurons, and second, cortical microcircuits are usually consisted of several hundreds of neurons, whereas the core receptor networks can have as few as four nodes [4].

We propose here a theoretical framework for cellular processing of time-varying signals in a vicinity of saddle-node (SN) bifurcation, where the information is encoded through metastable state generated via ghost attractor, and interpreted using phase-space trajectories. In contrast to networks with short-term memory, we demonstrate that the dynamical transient memory emerging due to this critical organization enables signal integration and optimal information processing even with two-component sensing systems. Using a single spiking leaky integrate-and-fire (LIF) neuron model [11] we demonstrate that these features are exhibited for a different type of intrinsic dynamics, and are thereby generic. Drawing a parallel to epidemic spreading on networks, we explain how such a critical behavior can take place on a molecular level of cell surface receptors. Using the example of the epidermal growth factor receptor (EGFR) system, we propose that a simple fluctuation-sensing mechanism could enable information processing capabilities to emerge in a self-organized manner, as a generic means to balance between plasticity and robustness in cellular sensing.

## II. DYNAMICAL INFORMATION PROCESSING AT CRITICALITY USING STATE-DEPENDENT TRAJECTORIES

Computational outcomes of systems that sense timevarying signals depend not only on the current stimulus, but also on memory of the previous sequence(s) in order to integrate the signal information [5]. The necessity for such a memory in receptor activity on the cell surface is even more pronounced to enable robust signaling response, since the majority of the receptors are rapidly internalized upon ligand binding and unidirectionally trafficked towards the cell interior where they are degraded [12, 13]. In broad range of biological systems, attractor networks are generically thought to underlie these memory functions [14–16]. A minimal cellular sensing network that accounts for memory in receptor activity (R_a_) is a two-component toggle-switch (Fig. 1a), where the double-negative feedback interaction [4, 17, 18] between the active receptor and an inactivating enzyme, a protein P_DNF,a_, follows the law of mass action:

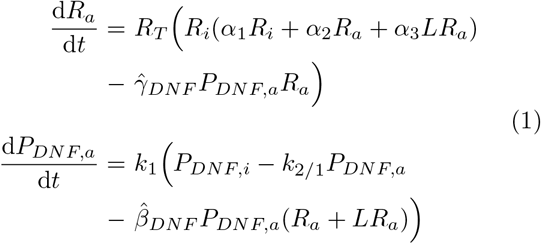

The system integrates the R_a_ activation- and the mutual inhibition mechanisms (Fig. 1a) that govern the protein state transitions between their active (*R*_*a*_, *P*_*DNF*,*a*_) and inactive (*R*_*i*_, *P*_*DNF*,*i*_) states. They are described in further detail in Appendix A with the corresponding parameters.

Bistability is exhibited between two saddle-node bifurcations for a broad range of the bifurcation parameter - the *P*_*DNF*,*T*_/ *R*_*T*_ concentration ratio - in absence of chemical stimulus input (Fig. 1b), and it is also maintained for a certain input range (Fig. 1c). The effective input for the cells in this case is the fraction of ligand-bound receptors that reflects the extracellular ligand concentration (Appendix A and [4]). Processing time-varying signals however, necessitates that the robustness that arises in attractor networks via bistability must be balanced with plasticity in the receptor responses in order to maintain sensitivity to upcoming stimuli.

To understand how this can occur in bistable systems, we studied qualitatively the dynamical R_a_-P_DNF,a_ behavior by analyzing how the phase-space trajectories evolve in relation to the changes in the geometry of the underlying phase space as a function of a pulsed stimulus. Generally, the relative positioning of the nullclines, which are determined by the system parameters, shapes the phase space geometry. In non-autonomous or input-driven systems, the underlying phase space can be altered either through its geometry (change in the positioning, shape and size of the attractors) or topology (change in the number or stability of the attractors) [19, 20]. We therefore also estimated the associated quasi-potential landscapes [21] (Figs. 1d to 1f, Appendix B and [19]). When starting from the valley of basal receptor activity in the double-well quasi-potential landscape characteristic for the bistable organization (Fig. 1b, green) in absence of stimulus (Fig. 1e middle), a topological phase space change that reflects the vanishing of this state occurs at a threshold signal concentration. This is manifested through a transition to the monostable state with high receptor activity (Fig. 1e, *i* → *ii* green transition). Upon signal removal, the reverse topological change leads to re-establishing of bistability (*ii* → *iii* green transition). However the trajectory remains in the occupied high activity steady state (green circle in Fig. 1e middle, and top). Thus, the first pulse will activate the receptors and this will hinder further responsiveness to upcoming stimuli due to the long-term memory that results from this stable attractor organization (Fig. 1e top, inset).

In contrast, organization in the monostable regime (Fig. 1b, blue) does not result in memory in receptor activity (Fig. 1d top and inset). The continuous and reversible repositioning of the single steady-state attractor that captivates the state trajectory through the induced changes in the phase space geometry (adding/removing stimulus: *i* → *ii*, *ii* → *iii*, blue transitions, Fig. 1d) in this case leads to receptor response that closely follows that of the input (Fig. 1d top, inset). This indicates that neither binor monostability can simultaneously account for plasticity and robustness in cellular responses to timevarying chemical cues.

For positioning in the vicinity of the saddle-node bifurcation point however (Fig. 1b, magenta), there is only one stable attractor the basal activity state (Fig. 1f middle). However, due to the closeness to the bifurcation point, the dynamics of the system here is qualitatively different than the remaining the monostable region. For a supra-threshold input pulse, the transition from the basal monostable - to the high activity monostable state (*i* → *ii* magenta transition) via the bistable region, determines robust activation of the receptor, whereas upon input removal, these consecutive topological transitions are reversed. During the reversal, there is a delay between establishing the single stable attractor (magenta state *iii*), and the system trajectory converging to it (magenta state *iv*), resulting in prolonged receptor activity before relaxation to basal level (Fig. 1f top). This delay causes a transient memory in receptor activity that does not hinder further responsiveness of the system (Fig. 1f top, inset).

The transient memory is a consequence of the critical dynamical behavior near the SN bifurcation. In this organization, the nullclines intersect only once, indicating a single low stable steady state. However, they are positioned very close to one another in the phase space area where the high steady state would be stable (compare Figs. 1f and 1e, middle), resulting in a quasi-potential landscape with a very shallow slope (Fig. 1f middle, inset). Thus, when the system transits back from the bistable to the monostable region, the remnant of the saddle-node that disappeared in this transition generates a metastable state that continues to capture the incoming trajectories (See Supplemental Video [22]). Such delayed dynamics referred to as a feature of a ghost attractor [23], has been previously shown in some driven dynamical systems such as ferroelectrics or semiconductor lasers [24]. In this organization, the system does not process information using the stable attractors associated with specific states (0-1 case), but rather the information is maintained via the transient memory and interpreted through the phase-space trajectories. The trajectories are in turn navigated through the dynamic quasi-potential landscape by the input-driven topological transitions.

**FIG. 1:**
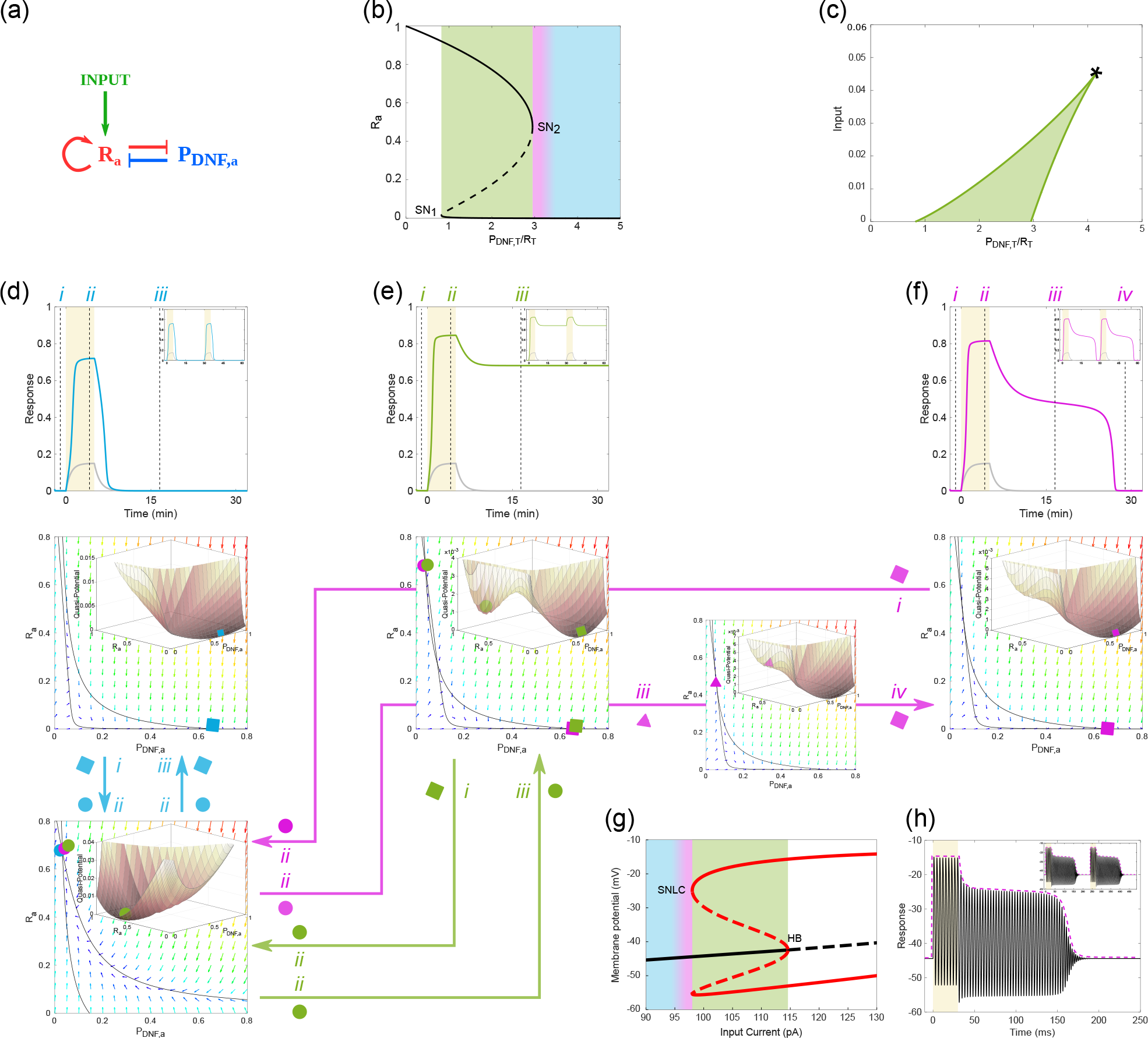
Plasticity and robustness to non-stationary cues emerges at criticality. (a) Diagram of a two-component toggleswitch between active receptors (R_a_) and the deactivating enzyme, protein P_DNF,a_. Input - fraction of ligand bound receptors (*LR*_*a*_). Molecular details described in Appendix A. (b) Bifurcation diagram of the R-P_DNF_ toggle-switch, depicting R_a_ response with respect to *P*_*DNF*,*T*_/*R*_*T*_, in absence of input. Shading: blue - monostable region, magenta - vicinity of the saddle-node (SN) bifurcation point, green - bistable region. (c) Two parameter (*LR*_*a*_, *P*_*DNF*,*T*_/*R*_*T*_) bifurcation diagram depicting the parameter space where bistability exists (green area). * - cusp bifurcation. (d) Top: Receptor response (blue) to single growth factor pulse (yellow) and subsequent input (LR_a_, grey) for positioning in the monostable regime (*P*_*DNF*,*T*_/*R*_*T*_ = 4.3). Inset: responsiveness to subsequent input pulses. Middle / Bottom: Phase space diagram and nullclines intersecting at the basal / high activity receptor steady state denoted with blue squares / circles respectively. Blue arrows: phase space transitions upon administering and removal of stimulus (*i* → *ii*, *ii* → *iii*, respective time points denoted in top). Insets: calculated quasi-potential landscapes. (e) Same as in d, only for positioning in the bistable regime (*P*_*DNF*,*T*_/*R*_*T*_ = 2.5). Green arrows: phase space transitions (*i* → *ii*, *ii* → *iii*). (f) Same as in d, for positioning at the critical transition between monostability and bistability (*P*_*DNF*,*T*_/*R*_*T*_ = 2.957). Magenta arrows: signal administration (*i* → *ii*) and removal (*ii* → *iii* → *iv*). The *iii* → *iv* transition and the associated phase space plot demonstrate the existence of a ghost attractor. (g) Bifurcation diagram of the LIF neuron model (Appendix C), depicting membrane voltage as a function of the input current (*I*_*app*_). HB: Hopf bifurcation, SNLC: saddle-node on limit cycle. Solid/dashed lines: stable/unstable steady state (black) and limit cycle (red). Shading same as in b. (h) Firing of a single LIF neuron during an input current increase (yellow) followed by transient memory after input resetting. Magenta: envelope of the neuronal response. Inset: Response upon consecutive inputs. Parameters: Appendix C.

This is a generic feature of systems organized at a critical proximity to a saddle-node bifurcation point. We demonstrate this on a system with different intrinsic dynamics - a single leaky integrate-and-fire (LIF) neuron model (Eq. (C1) in Appendix C). Bifurcation analysis showed that in this case, bistability between a resting and a spiking state is marked by a saddle-node on a limit cycle (SNLC) bifurcation (Fig. 1g). When organized in the vicinity of the SNLC, a continuous supra-threshold current input induces repetitive neuronal firing, which is transiently maintained after input resetting, due to the presence of the ghost of the SNLC (Fig. 1h). The envelope of the neuronal pulsing activity closely resembles the receptor activity profile at criticality (compare Fig. 1h and 1f top). Similar neuronal behavior where switching from a low-activity (baseline) state into a stimulus-selective activity state that is maintained throughout a delayed period after signal removal has been experimentally related to working memory processes in the large-scale neuronal networks in primate prefrontal cortex [25, 26]. It has also been demonstrated that stable attractor networks, as well as feedforward chain and random chaotic networks, cannot account for both stable coding as well as strong temporal dynamics of the neuronal population during working memory [26]. Thus, the critical behavior emerging for organization in the vicinity of the saddle-node bifurcation, although here simplified for a single neuron, could possibly provide the underling dynamical mechanism that supports working memory activity in the prefrontal cortex.

## III. TRANSIENT MOLECULAR MEMORY INTERPRETED THOUGH CRITICAL EPIDEMIC SPREADING

To understand how transient memory can be realized on a molecular level, we studied how transient receptor activity can be generated and maintained in absence of stimulus using single molecule reaction-diffusion simulation framework on a two-dimensional surface (Appendix D). Microscopic, or single-molecule receptor dynamics can be regarded as analogous to the susceptible-infected-susceptible (SIS) epidemic spreading system [27, 28]. In this analogy, single receptor molecules are susceptible to activation, equivalently to the S-state of an agent in the SIS model, and once active they can then propagate their state via direct contact (infected state of the SIS model). Here, receptor molecules can become susceptible again by interacting with active P molecules. The basic reproduction number *R*_0_ that determines the transmission potential of an infection [29] corresponds to the average number of newly ‘infected’ molecules by a single active receptor molecule in the course of its active lifetime, i.e. before its deactivation. If *R*_0_ < 1, the overall activity in the system decays (Fig. 2a, top), whereas if *R*_0_ > 1, the system exhibits supercritical behavior and the activity propagates in an epidemic-like branching fashion (Fig. 2a, bottom).

To relate the specific *R*_0_ realization when crossing the saddle-node bifurcations, we derived analytically the dependence of *R*_0_ to the main parameter that determines the positioning of the system in the vicinity of the SN bifurcation. For simplicity, we use *γ*_*DNF*_ as a bifurcation parameter, which is proportional to the specific reactivity of P_DNF,a_ to R_a_ (*γ*_*DNF*_ ∝ *P*_*DNF*,*T*_/*R*_*T*_, Fig. 1b). The activation transmission potential in every point of the phase space can be described as *R*_0_ ≡ *α*_2_*R*_*T*_ (1 − *R*_*a*_)/(*γ*_*DNF*_ *P*_*DNF*,*T*_ *P*_*DNF*,*a*_), where *α*_2_*R*_*T*_ (1 − *R*_*a*_) and 1/(*γ*_*DNF*_ *P*_*DNF*,*T*_ *P*_*DNF*,*a*_) are the average molecular activation rate and lifetime, respectively (kinetic parameters given in Appendix D). When initiated at the basal state, hence with a fully susceptible population (*R*_*a*_ = 0, *P*_*DNF,a*_ = *k*_1_/(*k*_1_ + *k*_2_), Appendix D), due to *R*_0_ = *α*_2_*R*_*T*_ (*k*_1_ + *k*_2_)/(*γ*_*DNF*_ *P*_*DNF*,*T*_ *k*_1_) ∝ 1/*γ*_*DNF*_ the critical threshold *R*_0_ = 1 is crossed at the *γ*_*DNF*_ value corresponding to SN1 (Fig. 2b bottom and middle; see Appendix D). For *γ*_*DNF*_ smaller than SN1, activity propagation is ensued as *R*_0_ > 1. However, once the system reaches the high activity steady state, *R*_0_ is maintained at 1 (Fig. 2b top and middle; Appendix D), because mass conservation limits the number of susceptible molecules. The system loses the transmission potential for *γ*_*DNF*_ values higher than SN2.

Equivalently to the macroscopic case (Figs. 1d to 1f), the phase space trajectories showed a clear attraction towards a single- (Fig. 2c top) or two coexisting attractors for different initial conditions (Fig. 2c middle) and different *γ*_*DNF*_ values. In the latter case, dynamical switching between the two steady states was observed. In the vicinity of SN2 however, the system exhibits critical behavior *R*_0_→1 (Fig. 2b, magenta region). The proximity of *R*_0_ to 1 together with the diffusion-induced spatial inhomogeneities in *P*_*DNF*,*T*_/*R*_*T*_ concentration can effectively increase the local transmission potential above the critical value 1, thereby generating local pockets of active receptor that transiently sustain and further propagate this state across the surface. This results in interchanging periods of inactivation and re-activation bursts around the ghost attractor state (e.g. green profile in Fig. 2c, lower-right panel), manifested as prolonged receptor activity before the system settles to basal state (Fig. 2c bottom equivalent to Fig. 1f).

**FIG. 2:**
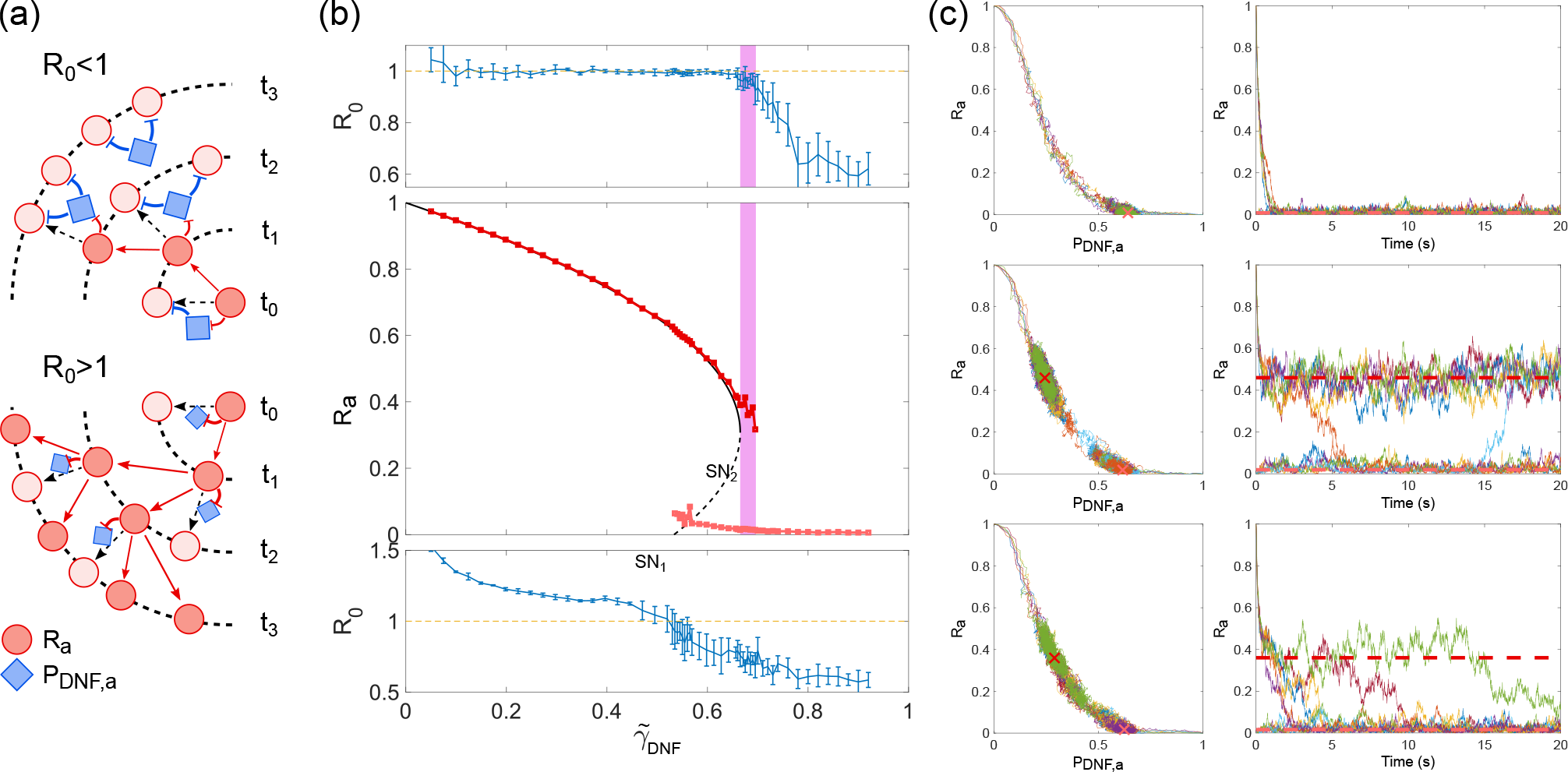
Molecular realization of transient memory. (a) Schematic representation of activity propagation in relation to microscopic single molecule activation/inactivation dynamics. Top: Diminished activity for *R*_0_ < 1. Bottom: Propagated activity for *R*_0_ > 1. (b) Basic reproduction number for varying *γ̃*_*DNF*_ values estimated by single-molecule reaction-diffusion simulations. Middle: Estimated high (dark red) and basal (light red) receptor activity states, black lines - estimated bifurcation profile; magenta - critical behavior. Top/bottom plots: *R*_0_ values around the higher (top) and basal (bottom) stable steady state. (c) Phase space R_a_-P_DNF,a_ (left column) and corresponding temporal profiles (right column) for organization in the three regimes. Trajectories represent the evolution of the average system state. Monostable regime (*γ̃*_*DNF*_ = 0.78, top), bistable regime (*γ̃*_*DNF*_ = 0.64, middle) and criticality (*γ̃*_*DNF*_ = 0.68, bottom). Red cross markers (left column) - estimated high and basal activity states; dashed lines (right column) - corresponding activity levels. Other parameters in Appendix D.

## IV. TRANSIENT VS. SHORT-TERM MEMORY: SIGNAL INTEGRATION AND OPTIMAL INFORMATION PROCESSING

Coming back to the macroscopic description of the system, the transient memory does not only enable plasticity and robustness in receptor responses, but it is also a unique manifestation of a dynamic memory. In other words, the total duration of the transient memory in receptor activity depends on the previous stimulus history, thus enabling signal integration (Fig. 3a, magenta). This feature arises from the metastability of the ghost state where the phase space trajectory gets transiently captured after stimulus removal, thus permitting responsiveness to subsequent signals. Equivalent behavior was observed also for the neuronal model (Fig. 3a, inset).

The stable attractor realization of a simple shortterm memory (reversible bistability) on the other hand emerges through the hysteresis of signal activation/deactivation levels (Fig. 3b). This memory is not manifested through prolonged temporal activation as for the transient memory (Fig. 3c, magenta), shown through step-wise input modulation (Fig. 3c, orange). This in turn excludes processing temporal dependencies that are inherent in time-varying signals, such that signal integration does not take place (Fig. 3a, orange). The other stable attractor realization, the long-term memory (irreversible bistability, Fig. 3c, green), does not allow responsiveness to upcoming signals to take place (Fig. 1e).

This crucial feature of dynamic memory is further complemented with additional information processing capabilities to enable optimal computation of time-varying signals. In the case of cell surface receptor networks for example, organization at criticality also endows cells with robust sensing via receptor activation in a switch-like manner (Fig. 3d), as a consequence of the topological phase space transitions (Fig. 1f, *i* → *ii*). Even more, the dynamic range of the response amplitude rapidly increases when transiting from the monostable towards the bistable regime, with a clear peak at the SN2 bifurcation (right to left, Fig. 3e). Using the single-molecule reaction-diffusion simulation framework, it can be additionally demonstrated that robustness to noise is also optimal at criticality: the probability for spurious activation in absence of stimulus highly increases in the bistable region, whereas in the vicinity of the SN2 bifurcation, this probability is close to zero (Fig. 3f). These results therefore show that critical organization is crucial for optimal information processing of time-varying signals.

**FIG. 3:**
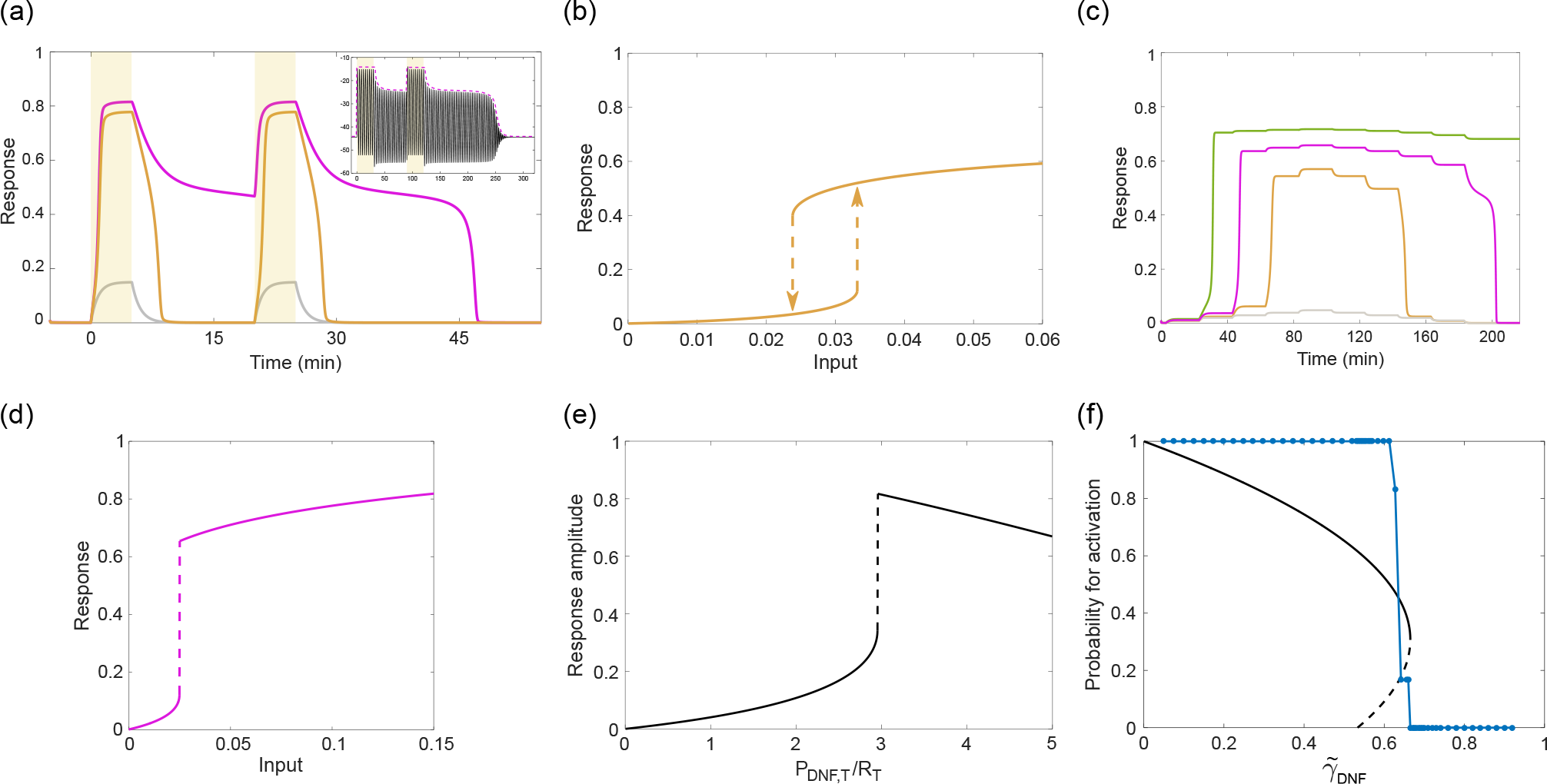
Signal integration and optimal information processing in the vicinity of a saddle-node bifurcation. (a) Receptor responsiveness to subsequent time-varying signals (yellow: ligand, grey: input/LR_a_) for positioning in reversible bistable (short-term memory, orange, *P*_*DNF*,*T*_/*R*_*T*_ = 3.5) and critical organization (transient memory, magenta). Inset: Signal integration (and response envelope, magenta) for critical positioning of the neuronal model. (b) Short-term memory (reversible bistability) reflected in the hysteresis of the activation/deactivation strength of the input. (c) Receptor responses for organization in long-term (green), transient (magenta) and short-term memory (orange) to step-wise input changes (grey). (d) Dose-response plot depicting receptor activity response (*LR*_*a*_ + *R*_*a*_) as a function of the fraction of ligand-bound receptors (*LR*_*a*_) for organization at the critical transition. (e) Dynamic range of receptor activation for input that activates the system (*LR*_*a*_ = 0.15), as a function of *P*_*DNF*,*T*_/*R*_*T*_. Other parameters as in Fig. 1. (f) Probability for spurious receptor activation (blue) overlaid with the estimated bifurcation profile (black), using the single-molecule simulation framework as in Fig. 2. Parameters: Appendices A and D.

## V. SELF-ORGANIZED POSITIONING AT CRITICALITY BY FLUCTUATION SENSING

We next investigated how such positioning can be realized for receptor networks. For this, we use the example of the proto-oncogenic epidermal growth factor receptor (EGFR), for which a critical organization between a monostable and bistable mode of operation was recently experimentally demonstrated [4]. The dynamics of EGF sensing in this case is regulated by a four-component spatially-distributed network, where the interactions of EGFR (R) with three specific membrane-associated enzymes - protein tyrosine phosphatases via a double negative (P_DNF_, PTPRG), a negative feedback (P_NF_, PTPN2) and a negative regulation (P_NR_, PTPRJ) are coupled via the vesicular trafficking of the receptor (Fig. 4a). Ligand-bound EGF receptors (LR) promote autocatalytic activation of ligandless receptors (red arrows) [12, 17, 18, 30], and thereby transfer information about the extracellular environment before they are internalized and degraded [12, 13]. The vesicular recycling on the other hand, brings the internalized and deactivated ligandless EGFR back to the plasma membrane, thereby establishing the EGF signal processing network. Numerical simulations using a two-compartment model that takes the trafficking-induced redistribution of receptors explicitly into account (Eq. (E1) in Appendix E) showed that input-induced decrease in plasma membrane receptor concentration rapidly shifts the operation of the system into the monostable regime. This results in decreasing dynamic range of the activation response amplitude and loss of transient memory of the recent stimuli (Fig. 4b, compare blue to magenta lines). Thus, the EGFR concentration at the membrane, previously a determining bifurcation parameter, must be dynamically maintained at the critical transition. Two-parameter bifurcation analysis showed how the positioning of the SN bifurcation point depends on the total receptor concentration *R*_*T*_ and its recycling rate constant *k*_*rec*_ (Fig. 4c).

**FIG. 4:**
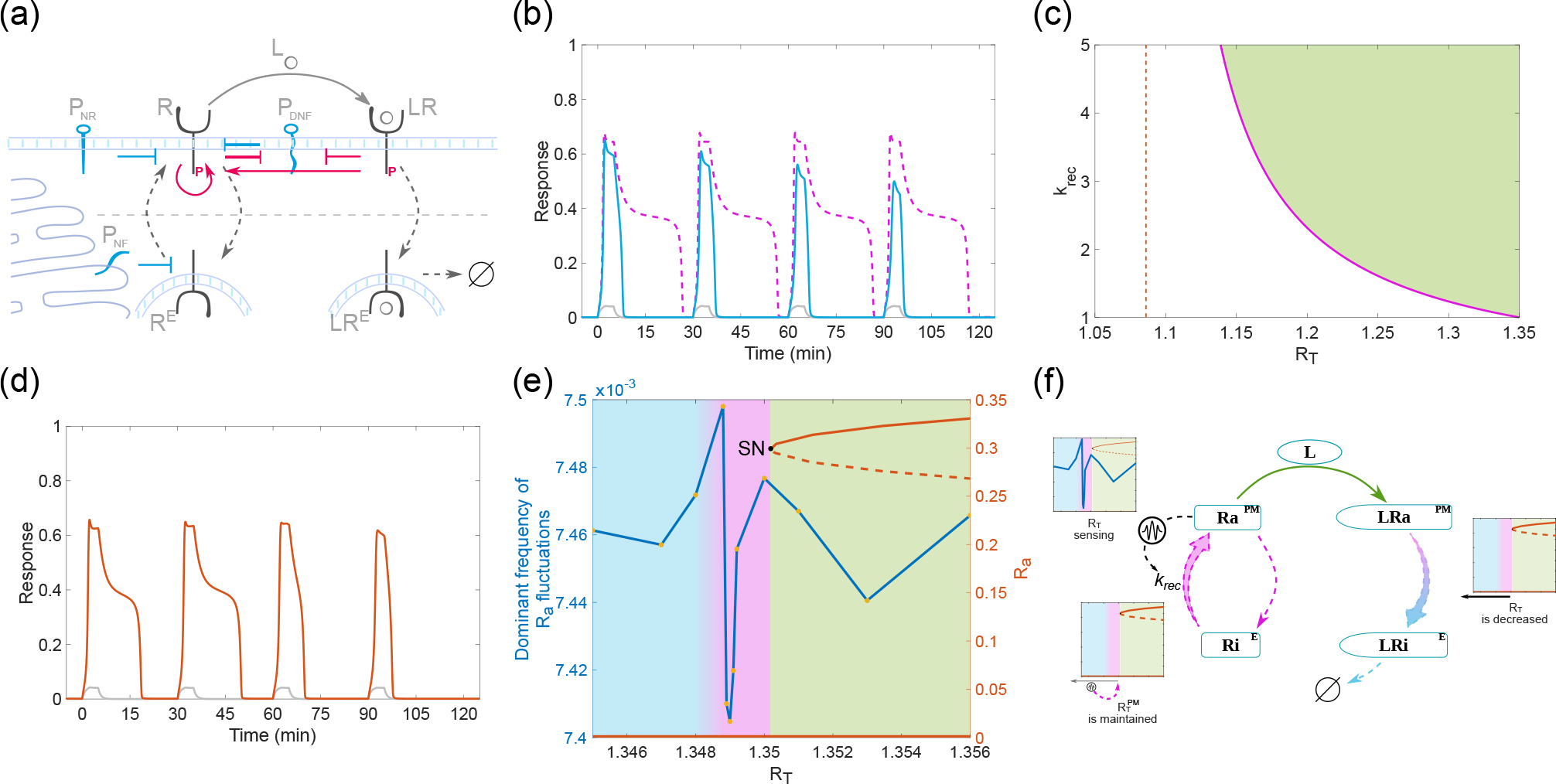
Dynamical mechanism for self-organization at criticality. (a) Schematic representation of the spatially-distributed EGFR network. At the plasma membrane, ligandless EGFR (R) is coupled to PTPRG (P_DNF_) in a double-negative feedback manner, and negatively regulated by PTPRJ (P_NR_). Activated receptors are then endocytosed in the perinuclear area (R^E^), deactivated by PTPN2 (P_NF_), and recycled back to the plasma membrane, which amounts to a spatially-established negative feedback. Ligand (L) binding converts R to ligand-bound species (LR) that are internalized (LR^E^) and subsequently degraded. LR promotes autocatalytic activation of R (red arrows). (b) Response of active (blue) and ligand-bound (grey) receptors at the cell surface, when degradation of ligand-bound receptors is explicitly considered (Appendix E), compared to the case without degradation (dashed magenta line). (c) Two-parameter (*k*_*rec*_, *R*_*T*_) bifurcation diagram depicting the positioning of the saddle-node bifurcation point SN2 (magenta line). Green shaded area: bistability region, red dashed line: asymptotic limit of the saddle-node positioning. (d) Dynamically maintained transient memory in receptor activity upon train of stimulus pulses. (e) Continuation plot of receptor activity (red) as a function of total receptor concentration and dominant frequency of fluctuations in basal receptor activity (blue) estimated for specific *R*_*T*_ values. Shading equivalent to Fig. 1b. (f) Schematic representation of a fluctuation-sensing and actuating system that dynamically poises receptor concentration at the plasma membrane in the vicinity of the saddle-node bifurcation point. Parameters: Appendix E.

Therefore, a self-organizing mechanism by which *k*_*rec*_ is up-regulated as a result of the decrease in total receptor concentration would effectively retain the system in the vicinity of the SN for several subsequent growth factor pulses (Fig. 4d). It should be noted however, that the SN positioning asymptotically approaches a minimal receptor concentration below which the receptor recycling rate can no longer sufficiently compensate for the loss of receptors from the membrane (Fig. 4c, dashed line). Additionally, the receptor recycling machinery may also impose an upper bound on *k*_*rec*_, further limiting the resetting capacity.

Such a dynamically-maintained organization would require a mechanism for sensing receptor abundance to estimate the divergence from the saddle-node bifurcation point, and an actuating mechanism to translate this positioning into a corresponding recycling rate. Information about the amount of receptors, especially in noisy settings, could be encoded in the fluctuations of the activity state. The temporal signature of these fluctuations depends on the alignment of the nullclines - hence on the positioning of the system in parameter space [8], which is in this case determined by the concentration of receptors. The dominant frequency in basal EGFR activity fluctuations that we estimated as an average of multiple stochastic activity profiles (Appendix G) was lowest around the SN bifurcation [24] (Fig. 4e). It is therefore possible to postulate an actuating molecular mechanism where cells use an effector protein downstream of EGFR that is embedded in a simple low-pass filter to up-regulate EGFR recycling when low frequency fluctuations of EGFR activity are present and thereby maintain positioning in the SN bifurcation vicinity (Fig. 4f). The previously identified characteristics of Akt [31–33], a serine-threonine kinase downstream of EGFR, qualify it as a candidate that could induce this prolonged maintenance of the system at the critical transition. However, once the system escapes the low-frequency valley around the SN and enters the monostable region, the cell will rely on EGFR synthesis to re-establish the organization at criticality in the long term. This example thus demonstrates that even a simple low-pass filter can help cells to maintain positioning at criticality from which optimal information processing capabilities in growth factor sensing emerge.

## VI. DISCUSSION

The physics of how cells perceive time-varying chemical signals is a fundamentally important question that dates back to the work of Berg and Purcell [34]. In the past decades, the limits of biochemical sensing have been explored using equilibrium and non-equilibrium descriptions of sensing through ligand binding/unbinding events to stationary receptors [34–39]. The ability of most receptors to detect low ligand concentrations however necessitates high affinity binding [40–42], limiting the detection of rapidly varying signals. Furthermore, the localization of many cell surface receptors is dynamically maintained through vesicular recycling, whereas upon ligand binding, unidirectional internalization and receptor degradation occurs [12, 13, 43, 44]. Although a rapid removal of ligand-bound receptors from the cell surface would in principle enable sensing of non-stationary cues via the remaining ligandless receptors, it would not allow for signal integration and robust signaling responses.

From the dynamical system formalism it follows that the topology of the receptor network determines the dynamical possibilities of the system. However, as we showed here, the optimal positioning in parameter space is what enables biochemical sensing of non-stationary environments via robust but plastic non-linear activity receptor responses. Such a critical self-organized behavior [45] resembles the one of the mammalian hearing system [46], flocks of birds [47] or biofilms [48].

We demonstrated that processing non-stationary inputs in real-time as ubiquitous during chemical sensing, necessitates information to be encoded through metastable states and interpreted using phase-space trajectories. The minimal requirements for such processing mechanism are met by the existence of a ghost of a saddle-node. This mechanism can be realized for various intrinsic dynamics and network sizes, as we have shown for two-component cell surface receptor network and single LIF neuron. In both cases, the dynamic transient memory that emerges from the critical organization allows for history-dependent signal integration and generally, optimized information processing - computational features that were previously only attributed to largescale neuronal networks [49–54].

Conceptually, the framework of information processing with state-dependent trajectories thus suggests that the notion of computation with stable attractors [55, 56], resembling the legacy of von Neumann and Turing [6], likely should be adapted for cellular processing of nonstationary signals. In this case, the dynamic transient memory and thereby related information processing features that emerge at criticality ensure optimal balance between stability and overall responsiveness to upcoming cues, as pervasive for systems that operate in a continuously changing environment.

## ACKNOWLEDGMENTS

The authors thank Philippe Bastiaens for numerous discussions and suggestions that were crucial for the development of this work, as well as for critically reading the manuscript.

A.K. conceptualized the study. A.S. performed most of the analytical derivations, numerical simulations and bifurcation analysis with help of A.P.N. and A.K. A.S. and A.K. interpreted the results and wrote the manuscript with help of A.P.N.

The authors declare no conflict of interests.

## APPENDIX

## Appendix A: Modeling the R-PDNF toggle-switch

With Eq. (1) we model a minimal network motif that exhibits bistability, a double-negative feedback, using law of mass action. Both, the receptor, as well as the deactivating enzyme have active (R_a_, P_DNF,a_) and inactive (R_i_, P_DNF,i_) states, and their state transition rates are described by the model equations. Therefore, mass is conserved in the system and the total protein concentrations of both species (*R*_*T*_, *P*_*DNF*,*T*_) are constant parameters. This allows *R*_*i*_ = 1 − *R*_*a*_ and *P*_*DNF*,*i*_ = 1 − *P*_*DNF*,*a*_, expressed as fractions from the total protein concentrations. Since the fraction of ligand-bound receptors (*LR*_*a*_) is mapped to the cell from the ligand concentration in the environment [4], it is considered as an input parameter in the system.

Autonomous, autocatalytic and ligand-bound-induced activation of ligandless R_i_ ensue from bimolecular interactions with distinct rate constants *α*_1−3_, respectively. Other parameters: *k*_2/1_ - P_DNF_ activation/inactivation rate constant ratio, *k*_1_ - kinetic constant that does not influence the steady state values of the system, 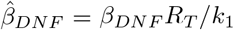 - receptor-induced regulation rate constant of P_DNF_, 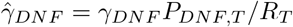 - specific reactivity of the enzyme towards the receptor, thus proportional to the local effective *P*_*DNF*,*T*_ */R*_*T*_ ratio. In the analysis we refer to changes of *P*_*DNF*,*T*_ */R*_*T*_ when numerically 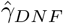 is varied.

For simulations with growth factor pulses in Figs. 1d to 1f and Fig. 3a, binding/unbinding of ligand to modulate *LR*_*a*_ was introduced. Thus, 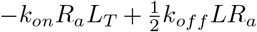, was added as additional term to the differential equation of R_a_, and the dynamics of LR_a_ was modeled with 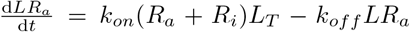. For the simulations in the Supplementary Video [22] a stochastic differential equation model was constructed from Eq. (1) by adding a multiplicative noise term *σX*_*i*_(1 − *X*_*i*_)d*W*_*t*_, where *σ* = 0.05, d*W*_*t*_ is the Brownian motion term and *X*_*i*_(1 − *X*_*i*_) is the state-dependent function for each variable *i* that accounts for mass conservation and normalization of the variables. The model was solved with Δ*t* = 0.01 using the Euler solver from the Financial Toolbox in MATLAB. The parameters corresponding to Figs. 1a to 1f and Figs. 3a to 3e are: *α*_1_ = 0.0017, *α*_2_ = 0.3, *α*_3_ = 1.0, 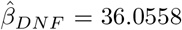, 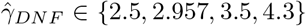 (bistability, memory, reversible bistability, monostability), *k*_1_ = 0.01, *k*_2/1_ = 0.5, *R*_*T*_ = 0.8, *k*_*on*_ = 0.003, *k*_*off*_ = 0.01668. The *L*_*T*_ amplitude during the pulse was set to produce 0.15 of *LR*_*a*_ in steady-state.

## Appendix B: Quasi-potential landscape computation

The numerical computation of the quasi-potential landscapes, corresponding to the phase space diagrams in Figs. 1d to 1f was conducted using an approach adapted from the one in [57]. Multiple trajectories were calculated starting from initial conditions distributed on a grid in the phase space. Rate of change in the quasi-potential (initiated arbitrarily to 0) was calculated by 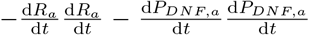 in every iteration and the quasi-potential was integrated by the ODE solver together with the system trajectory integration. Quasi-potentials of trajectories converging to the same attractor were aligned to match at the steady-state level. Quasi-potentials of different attractors were subsequently aligned at the initial points of the neighboring trajectories that converge to the different respective attractors, i.e. at the separatrix points. Additionally, neighboring separatrix pairs were weighted by the angle between their derivatives (*θ*), according to the formula: 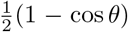. This gives greater weight to diverging pairs, effectively aligning the separatrix quasi-potential values at the saddle steady state point. The quasi-potential landscape at every point in phase space was finally estimated by interpolation from the aligned quasi-potential values of all of the trajectory points.

## Appendix C: Model of a leaky integrate-and-fire (LIF) neuron

We used a well-established model of a LIF neuron [11], described with the following set of equations:

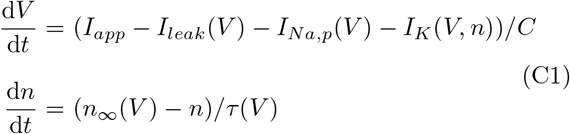

where *V* is the membrane potential and *n* - activation variable for K^+^. *I*_*app*_ is the input current, *I*_*leak*_ (*V*) = *gl*(*V* − *E*_*l*_) - Ohmic leak current, *I*_*Na*,*p*_(*V*) = *g*_*Na*_*m*_*∞*_(*V*)(*V* − *E*_*Na*_) - voltage-gated persistent Na^+^ current and *I*_*K*_ (*V*, *n*) = *g*_*K*_*n*(*V* − *E*_*K*_) - voltage-gated persistent K^+^ current. *m*_*∞*_(*V*) = 1/(1 + exp{(*V*_1/2,*m*_ − *V*)*/k*_*m*_}) and *n*_*∞*_(*V*) = 1/(1 + exp{(*V*_1/2,*n*_ − *V*)*/k*_*n*_}) can be approximated by the Boltzmann function. *τ* (*V*) = 1. Parameters: *g*_*l*_ = 1, *E*_*l*_ = −78, *g*_*Na*_ = 2.7, *E*_*Na*_ = 60, *g*_*K*_ = 4, *E*_*K*_ = −90, *C* = 1, *V*_1/2,*n*_ = −45, *k*_*n*_ = 5, *V*_1/2,*m*_ = −30, *k*_*m*_ = 7.

## Appendix D: Estimation of the basic reproduction number using single-molecule reaction diffusion framework

We considered a two-dimensional domain representing the plasma membrane containing reacting and diffusing single molecules. The spatial coordinates of the molecules were updated using Brownian dynamics and time was discretized to intervals of length Δ*t*. First-order unimolecular reactions occur spontaneously with proba-bility *k̃*Δ*t*, where *k̃* is the intrinsic reaction rate constant. Second-order bimolecular reactions on the other hand are modeled using the Doi method [58], following the Smoluchowski single-particle framework for describing diffusion influenced reactions [59]. An interaction takes place between two molecules that have diffused within a proximity distance *σ* of each other, and a reaction ensues with a probability *g̃*Δ*t*, where *g̃* is the microscopic bimolecular reaction rate constant. *σ* is of order of the molecule radius. In the rare event that a substrate molecule is in proximity of *n* > 1 other enzyme molecules, reaction takes place with probability 1 − (1 − *g̃*Δ*t*)^*n*^, assuming any of the enzyme molecules can affect the state of the given substrate molecule. Formation of the product proceeds immediately upon successful bimolecular enzyme-substrate interaction, i.e. the state of the substrate molecule is directly changed. To model the interactions between R and P_DNF_ (Fig. 1a), we assumed that both particles diffuse across the 2D domain with equal diffusion rates 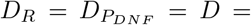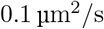. The interaction radius *σ* was set to 2*ρ*, where *ρ* = 10 nm is the molecule radius. 500 receptor molecules and 700 *P*_*DNF*_ molecules were randomly deployed on a 5 μm × 5 μm surface and allowed to diffuse using Brownian dynamics with periodic boundary conditions. Δ*t* was set to 1 × 10^−3^ s to ensure that 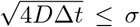, i.e. any interaction between two molecules that come in proximity is detected, and also to ensure that no reaction probability is greater than one. The state transitions of R and P_DNF_ occur in accordance with the macroscopic description - Eq. (1), in absence of external input (*LR*_*a*_ = 0). The microscopic rate constants are therefore proportional to the ones in our main ODE model: *α̃*_1_ = 0.0017/(*σ*^2^*π*), *α̃*_2_ = 0.3/(*σ*^2^*π*), *β̃*_*DN F*_ = 0.360558/(*σ*^2^*π*), *k̃*_1_ = 0.005/Δ*t*, *k̃*_2_ = 0.0025/Δ*t*. They were set to produce faster kinetics, due to numerical and data storage constraints. By analogy to the macroscopic bifurcation analysis (*γ̃*_*DN F*_ ∝ *P*_*DNF*_ */R*_*T*_), *γ̃*_*DN F*_ was varied between 0 and 0.92/(*σ*^2^*π*) (*P*_*DNF*,*T*_ and *R*_*T*_ were kept constant) to modulate the specific reactivity of P_DNF,a_ towards R_a_. To calculate the basic reproduction number *R*_0_, the number of substrate receptor molecules *R*_0,*j*_ (*t*) that each R_a,j_ molecule successfully activated within its activation lifetime was recorded, after that molecule has been previously activated itself at time *t*. *R*_0_ was calculated as an average of these figures within a certain time interval. Theoretically *R*_0_ depends on the probability of activating a susceptible receptor molecule and the duration of the activity lifetime, in analogy to the SIS epidemic spreading model. Neglecting the effect from autonomous activation of R (with low rate *α*_1_), we rive at the following definition 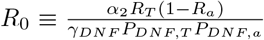. From Eq. (1) it is straightforward to determine the basal activity stable steady state as *R*_*a*_ = 0, 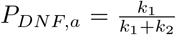, and thus 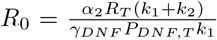. By employing linear stability analysis we could determine that the condition for stability of this steady state is indeed *R*_0_ < 1. On the other hand, it follows readily from the first equation that *R*_0_ = 1 around the high activity stable steady state, where *R*_0_ ≠ 0. There we could also find that 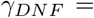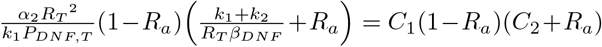, thus there is approximately a quadratic dependence between *γ̃*_*DNF*_ and *R*_*a*_. This form was used in Fig. 2b, middle, to estimate the bifurcation profile: *C*_1_ = 1.388, *C*_2_ = 0.385, which translates to *γ̃*_*DNF*,*SN*1_ = 0.5344, *γ̃*_*DNF*,*SN*2_ = 0.66. To estimate the extensively occupied high and basal receptor activity states from the trajectories in R_a_-P_DNF,a_ phase space (Fig. 2c), Gaussian mixture distribution was fitted to the data with two components and data points were pruned iteratively with 90th percentile cut-off until convergence. The trajectories, as well as temporal profiles on Fig. 2c were visualized with downsampling to every fifth frame due to size constraints, without loss of visual quality.

## Appendix E: Compartmental model of spatial-temporal EGFR activation regulation

The experimentally derived EGFR-PTP network [4] was implemented using a two-compartmental model that includes explicitly the vesicular trafficking between the plasma membrane and the endosomal compartments (Fig. 4a), as described using the following system of ODEs:

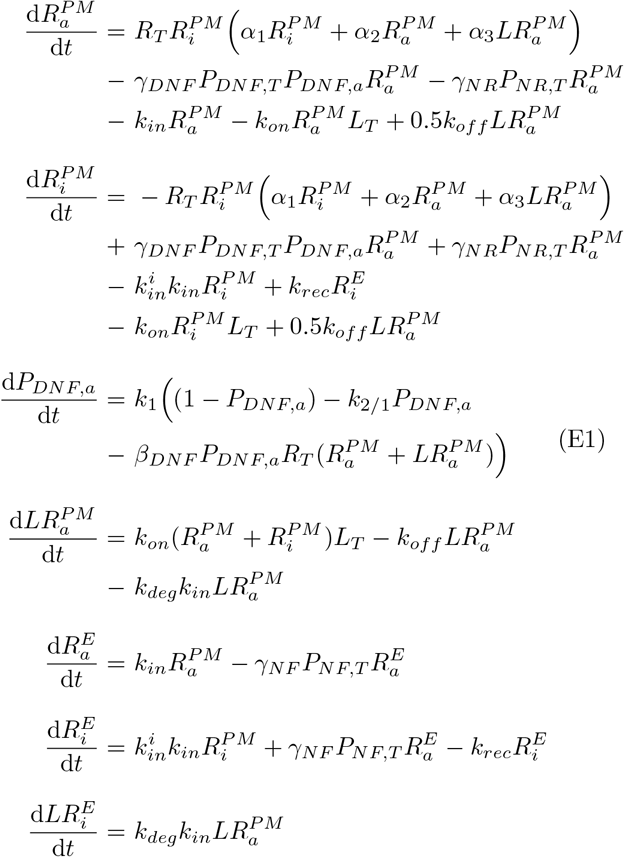

P_DNF_ (PTPRG), P_NF_ (PTPN2), P_NR_ (PTPRJ) represent the major protein tyrosine phosphatases that regulate EGFR (R, LR) activity, 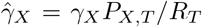 - specific reactivity that each P_X_ ∈ {P_DNF_ (PTPRG), P_NF_ (PTPN2), P_NR_ (PTPRJ)} has towards EGFR and is therefore proportional to local effective *P*_*X*,*T*_ */R*_*T*_ ratio. *k*_*in*_, *k*_*rec*_, *k*_*deg*_ denote receptor internalization, recycling and degradation rate constants, respectively, *i*,*a* - inactive and active species, *E*, *PM* - endosomal and plasma membrane species. For the self-organizing criticality model (Fig. 4d), the dynamically maintained 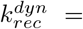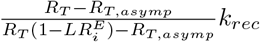, where *R*_*T*, *asymp*_ = 1.086 is the lower bound asymptotic value of *R*_*T*_ in dependence to *k*_*rec*_ (dashed line, Fig. 4c). Saturation level of 2.5 for the multiplier term is also assumed, beyond which the recycling rate can no longer increase. Parameters: 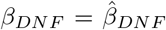, *γ*_*DNF*_ = 3.0, *γ*_*NF*_ = 3.0, *γ*_*NR*_ = 0.001, *k*_*in*_ = 0.02, *k*_*rec*_ = 0.042, 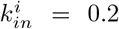, *k*_*deg*_ = 0.2, *R*_*T*_ = *P*_*DNF*,*T*_ = *P*_*NF*,*T*_ = *P*_*NR*,*T*_ = 1.0. Other parameters are the same as in Fig. 1.

## Appendix F: Model calibration

The parameters in the model (Fig. 1a and Fig. 4) were described in [4] and calibrated with model-based fits of the single cell dose-response data from where the topology of the sensing network (Fig. 4a) was derived. We convert from dimensionless time to minutes by equating the EGFR phosphorylation kinetics and duration in the simulations using the kinetic parameters to the experimental values in [4]. The parameters for the microscopic dynamics in the single-molecule reaction diffusion simulations were set to scale the macroscopic ODE parameters and set to produce faster kinetics due to numerical reasons, as described in the corresponding section.

## Appendix G: Stochastic simulations

To model the contribution of random fluctuations in the activity dynamics of the network constituents, a multiplicative noise term was added to the first equation in the ODE system (Eq. (E1)): 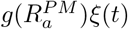. *ξ*(*t*) is a Gaussian white noise with zero mean and temporal correlation: 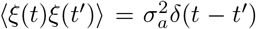, where *δ*(*t* − *t*′) is the Dirac delta function and 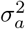 is a constant that characterizes the noise intensity. The multiplicative noise term is interpreted according to Stratonovich [60], as a stochastic interpretation for a realistic noise with small temporal autocorrelation [61]. This noise term can incorporate both intrinsic and extrinsic sources [62]. We establish the function 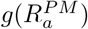 by means of simple approximation, assuming that the relative fluctuations scale is the inverse square-root of the amount of active protein. Such scaling is generic for many stochastic processes (e.g. Poisson processes or birth-death processes) and provides means to investigate the implications of fluctuations on the dynamics of biochemical networks in general [62]. To avoid negative values of the protein concentrations due to the introduced stochasticity in the system, a custom-made adaptive step size algorithm [63] employed to Euler integration scheme was developed in C. *N* = 50 realization of stochastic time series were simulated with 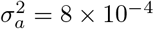 and for each of them the dominant frequency in basal receptor activity was extracted by computing the dominant mode of the wavelet power spectrum using the WaveletComp package in R [64]. The average dominant frequency from this realizations is plotted in Fig. 4e.

## Appendix H

The numerical bifurcation analysis was performed using the XPP/AUTO software [65].

All simulation except where explicitly noted were performed using custom-made code in MATLAB (MATLAB and Statistics Toolbox Release R2018a, The MathWorks, Inc., Natick, Massachusetts, United States).

All data and code used in this manuscript are available from the corresponding author upon reasonable request.

