## Supplemental Material for "Optimal biochemical information processing at criticality"

**Supplemental Video 1**. **Transient memory in receptor activity via a "ghost" attractor.** Phase space transitions and a trajectory (left) depicting receptor responsiveness (right, green – receptor activity, grey – fraction of ligand-bound receptors) to single stimulus pulse (yellow) for positioning at criticality. Orange circles - stable steady states, blue circle - unstable steady state, dashed line - separatrix. The results depict stochastic realization of the model equations (1) including ligand binding dynamics (see Appendix A).
